# Cilia-independent requirements for Ccdc103 promoting proliferation and migration of myeloid cells

**DOI:** 10.1101/2020.11.24.397026

**Authors:** Lauren G. Falkenberg, Sarah A. Beckman, Padmapriyadarshini Ravisankar, Tracy E. Dohn, Joshua S. Waxman

## Abstract

The pathology of primary ciliary dyskinesia (PCD) is predominantly attributed to impairment of motile cilia. However, PCD patients also have perplexing functional defects in myeloid cells, which lack motile cilia. Here, we show Coiled-coiled domain containing protein 103 (CCDC103), mutations in which underlie PCD, is required for the proliferation and directed migration of myeloid cells. CCDC103 co-localizes with cytoplasmic microtubules in human myeloid cells. Zebrafish *ccdc103*/*schmalhans* (*smh*) mutants have reduced macrophage and neutrophil proliferation, rounded cell morphology, and an inability to migrate efficiently to the site of sterile wounds. Furthermore, we demonstrate that direct interactions between CCDC103 and Sperm associated antigen 6 (SPAG6), which also promotes microtubule stability, are abrogated by CCDC103 mutations from PCD patients, and that *spag6* zebrafish mutants recapitulate the myeloid defects of *smh* mutants. In summary, we have illuminated a mechanism, independent of motile cilia, to explain functional defects in myeloid cells from PCD patients.

**Summary Statement:** We show Ccdc103 regulates myeloid migration and proliferation independent of cilia in zebrafish and that mutations in CCDC103 that cause primary ciliary dyskinesia abrogate interactions with the microtubule-stabilizing protein SPAG6.

## Introduction

Impairment of motile cilia results in primary ciliary dyskinesia (PCD), a disease found in 1:10,00-40,000 live births (Damseh *et al*., 2017). PCD is a genetically and phenotypically heterogeneous disorder that often presents with organ laterality randomization, infertility, and recurrent respiratory infections due to impaired mucociliary clearance (Leigh *et al*., 2009; Horani *et al*., 2016; Shapiro *et al*., 2018). While most of the phenotypic hallmarks of PCD are attributable to organs that require motile cilia, it has also been hypothesized that an additional primary immune dysfunction might contribute to the chronic lung infections seen in PCD patients. Intriguingly, studies spanning multiple decades, most of which antedate any identification of genetic lesions in PCD patients, report that myeloid cells isolated from PCD patients have functional defects, including disrupted migration in response to chemotactic stimuli and dysregulated expression of surface receptors (Afzelius et al., 1980; Cockx et al., 2018; Englander and Malech, 1981; Valerius et al., 1983; Walter et al., 1990). Additionally, ultrastructural analysis coincident with these neutrophil migration studies showed disruption of cytoplasmic microtubules, including altered number and location of centrioles (Valerius et al., 1983). These observations, though made in multiple independent cohorts of patients, which likely contained different causative mutations, have never been mechanistically explained. Thus, understanding the molecular basis for defects in PCD patient myeloid cells may provide critical insights into this disease process and open new avenues for potential therapies to improve outcomes for what is a life-long and often debilitating condition.

Presently, mutations in more than 40 different gene products have been implicated in PCD (Horani *et al*., 2016). The genes most commonly mutated in this disorder include light, intermediate, and heavy chain subunits of axonemal dynein and its required chaperones/assembly factors, which are required for ciliary motility, including Dynein axonemal assembly factor 1 (DNAAF1), and microtubule binding proteins such as sperm-associated antigen 1 (SPAG1). One of these axonemal dynein assembly factors is Coiled coiled domain-containing protein 103 (CCDC103). Mutations in *CCDC103* underlie approximately 4% of all PCD cases, but in certain geographic subpopulations where PCD is more prevalent it has been shown to be responsible for ∼20% of cases (Shoemark *et al*., 2018). CCDC103 is a small 242 amino acid (AA) protein that consists of a central RPAP3_C domain flanked by N- and C-terminal coiled coils (King and Patel-King, 2020). It is critical for the proper docking and assembly of the outer dynein arms which facilitate ciliary motion, but it is also found localized throughout the cytoplasm of both ciliated and unciliated cells (Panizzi *et al*., 2012). *In vitro* studies have shown that CCDC103 forms self-organizing oligomers, and that it binds periodically to cytoplasmic microtubules and can facilitate the stability of assembled microtubules (King and Patel-King, 2015; King and Patel-King, 2020). Despite these studies, if CCDC103 has cilia-independent requirements within the cytoplasm *in vivo* remains unknown.

Here, we show that *Ccdc103* has conserved expression in vertebrate myeloid lineages, including primitive macrophages and neutrophils, and localizes with cytoplasmic dynein (DYNH1C1) on microtubules within their cytoplasm. Using zebrafish *ccdc103*/*schmalhans (smh)* mutants, an established model for PCD, we find their myeloid cells have decreased proliferation, disrupted directed migration to sterile wound sites, and an abnormal spherical morphology, findings which are consistent with a loss of cytoplasmic microtubule stability. Interestingly, we identified Sperm-associated antigen 6 (SPAG6), which promotes microtubules stability and is associated with proper proliferation and migration in multiple cells types, as a novel CCDC103-binding partner. Patient mutations in CCDC103 abrogate interactions with SPAG6, while engineered zebrafish *spag6* mutants recapitulated functional defects in myeloid cells found in *smh* mutants. Taken together, our study is the first to identify motile cilia-independent roles of CCDC103 and illuminate a mechanism underlying unexplained functional defects in PCD patient myeloid cells, which may open new avenues to improve outcomes for these patients.

## Results

### *Ccdc103* is expressed in zebrafish myeloid progenitor cells

In an *in situ* hybridization (ISH) screen for novel genes expressed in the anterior lateral plate mesoderm (ALPM), we observed that *ccdc103* is expressed lateral to the developing head in 17 somite stage (ss) embryos (Fig. 1A,B). This region in zebrafish gives rise to an anterior population of myeloid progenitors and the expression was surprising given Ccdc103’s role in motile cilia (Panizzi *et al*., 2012; Austin-Tse *et al*., 2013; Casey *et al*., 2015). As previously reported (Panizzi *et al*., 2012), *ccdc103* was also highly expressed in the pronephros (Fig. 1A), an organ which requires motile cilia. As there is some heterogeneity in the progenitor cell types in the ALPM (Gering *et al*., 2003), we initially determined whether *ccdc103* is expressed in myeloid progenitors by co-injecting mRNA encoding the master hematopoietic regulators *scl* and *lmo2*. We found that the anterior *ccdc103* expression domain was expanded in injected embryos (Fig. 1C,D). RT-PCR for *ccdc103* in flow-sorted *spi1b:EGFP+* cells from 17ss embryos further supported its expression in myeloid progenitors (Fig. 1E). Additionally, RT-qPCR performed on cDNA from whole zebrafish embryos injected with the pro-myeloid pioneer factor *spi1b* showed a significant increase in *ccdc103* expression compared to controls along with a corresponding decrease in *gata1* expression, denoting a shift away from an erythroid fate (Fig. 1F) (Ward *et al*., 2003). Furthermore, RT-PCR for *CCDC103* supported it is expressed in human myeloid leukemia lines, including the promyelocytic leukemia HL-60 cell line, whole-blood derived CD34+/CD38-cells, and whole cord blood (Fig. 1G and Fig. S1). Thus, our data show that *Ccdc103* has previously unrecognized conserved expression in zebrafish and human myeloid cells.

**Figure 1.**
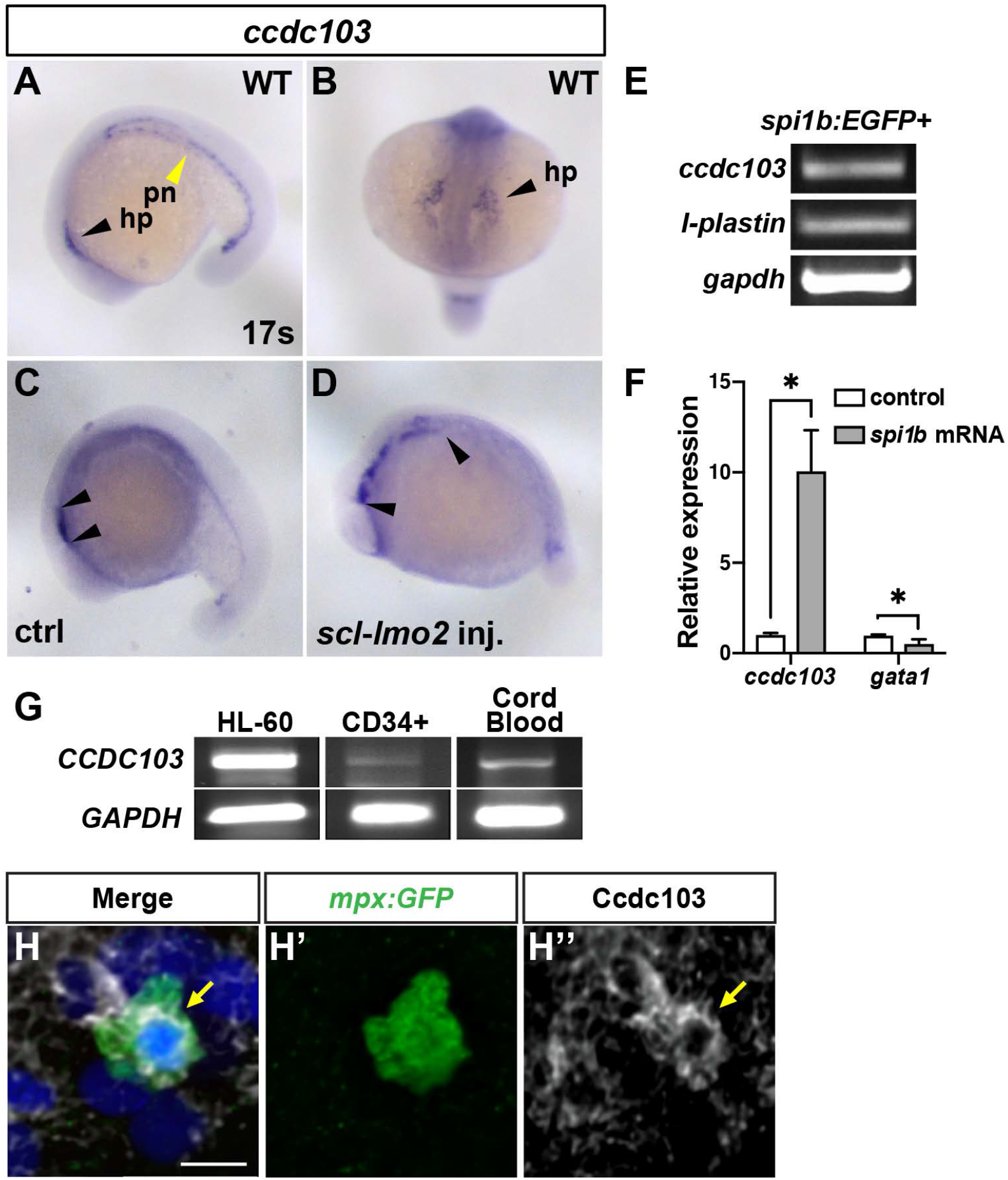
*Ccdc103* is expressed myeloid cells. **(A**,**B)** *Ccdc103* expression at 17ss WT embryo. hp - hematovascular progenitors (black arrowheads), pn – pronephros (yellow arrowhead). **(C**,**D)** The anterior domain of *ccdc103* expression (black arrowheads) is expanded in embryos co-injected with *scl* and *lmo2* mRNAs. In A,C, and D, images are lateral views with anterior left. In B, image is a dorsal view. **(E)** RT-PCR for *ccdc103, l-plastin* (+control), and *gapdh* performed on cDNA isolated from FACS-sorted *spi1b:EGFP*+ zebrafish myeloid progenitor cells. **(F)** RT-qPCR for *ccdc103* and *gata1* performed on cDNA isolated from embryos injected with *spi1b* mRNA. **(G)** RT-PCR for human *CCDC103* was done with cDNA isolated from HL-60 cells, CD34+/CD38-HSCs, and cord blood. **(H-H’’)** IHC for Ccdc103 (arrows), GFP, and DAPI in *mpx:GFP+* zebrafish neutrophil at 24 hpf. Scale bars: 10 μm.

Using immunohistochemistry (IHC) with a pan-Ccdc103 antibody, we also confirmed that Ccdc103 is present in primitive zebrafish *spi1b:EGFP+* myeloid cells where it appeared as puncta that were absent in *smh* embryos (Fig. S2 and Fig. S3). However, in *mpx:GFP+* neutrophils it had a perinuclear localization (Fig. 1H-H’’). To examine CCDC103 localization in human myeloid cells, we performed immunostaining on HL-60 cells, as well as HL-60-derived macrophage (MP)-like and neutrophil-like cells (Breitman, Selonick and Collins, 1980; Millius and Weiner, 2010). In the HL-60 myeloid progenitor cells, we observed CCDC103 localized in a larger, more sparse puncta, while in HL-60-derived neutrophils and macrophages CCDC103 had smaller punctate and a more diffuse distribution throughout the cells (Fig. 2A-B, 2E-F, 2I-J). Interestingly, the CCDC103 puncta in all these cells appeared to associate closely with the cytoplasmic microtubule network. Furthermore, in HL-60 myeloid progenitors, CCDC103 and Cytoplasmic dynein heavy chain 1 (DYNC1H1) aggregates were concentrated at putative microtubule organizing centers (MTOCs), as revealed with *α*-Tubulin (TUBA) (Fig. 2A-D), which was a consistent finding with zebrafish myeloid cells (Fig. 1H-H’). Though previous work has indicated myeloid cells are not ciliated (Finetti *et al*., 2009; Yuan and Sun, 2013; McClure-Begley and Klymkowsky, 2017) and there was no evidence of any cilia in the previous imaging, we performed staining for acetylated (K40) TUBA, which marks cilia. However, the acetylated TUBA was predominantly localized to putative MTOC and there was no indication of cilia (Fig. S4). In the differentiated HL-60 cells, CCDC103 was more broadly associated with the microtubule network and DYNC1H1, with the puncta often appearing to sit on top of the microtubules (Fig. 2E-L). Thus, we find that CCDC103 has conserved expression and co-localizes with cytoplasmic microtubules in vertebrate myeloid cells.

**Figure 2.**
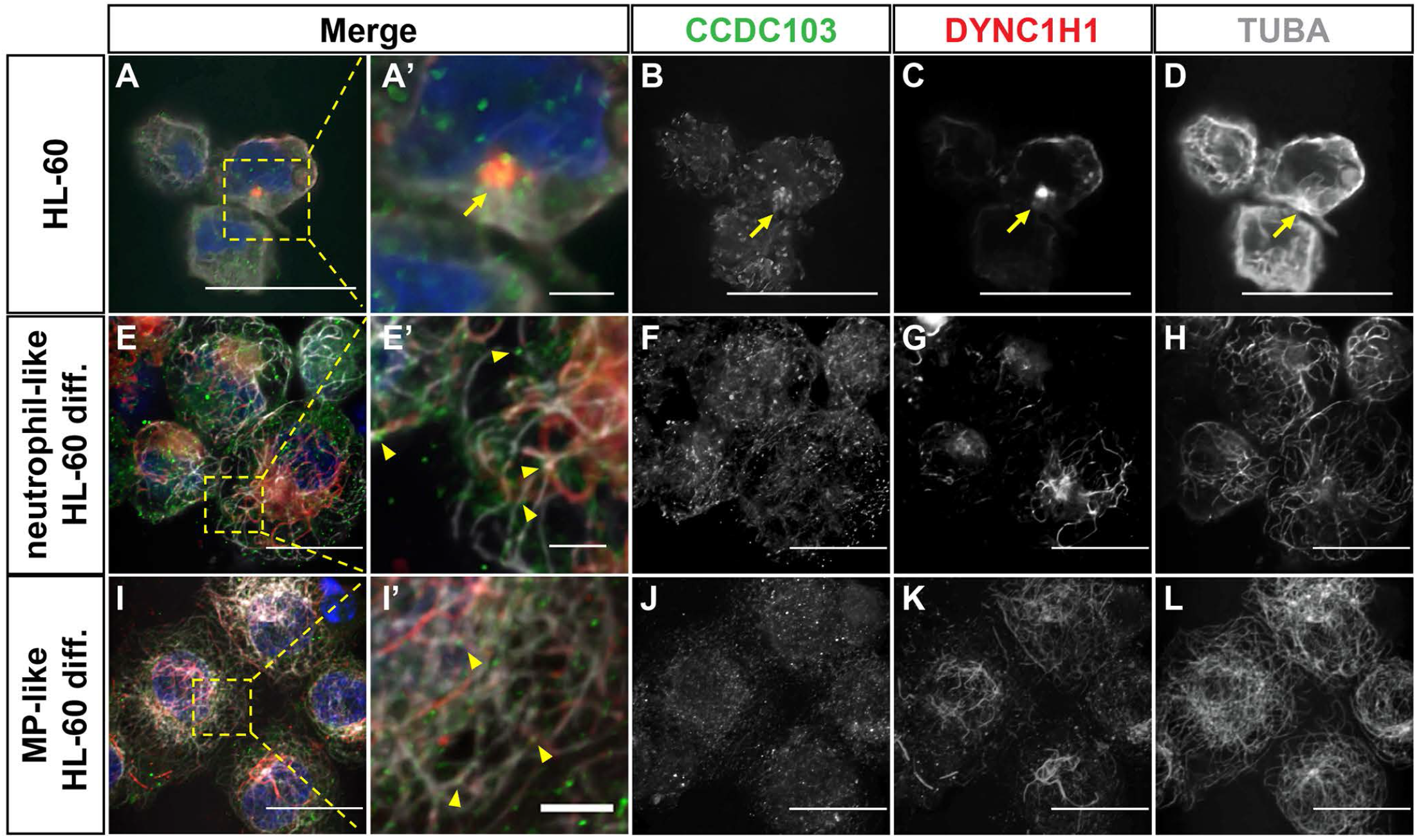
CCDC103 co-localizes with TUBA and DYNC1H1. **(A-D)** IHC for CCDC103, DYNC1H1, and TUBA in HL-60 cells. Yellow arrows - microtubule organizing center. **(E-H)** HL-60-derived neutrophil-like cells. **(I-L)** HL-60-derived macrophage-like cells. In E-L, yellow arrowheads - regions of colocalization of CCDC103 and TUBA in neutrophil-like and macrophage-like cells. Scale bars (A-L): 10 μm, Scale bars (A’,E’,I’): 2 μm.

### Myeloid cells in *smh* mutants have reduced proliferation

The zebrafish *smh* mutant has ciliary paralysis analogous to that seen in PCD patients and results from a point mutation that causes premature truncation of Ccdc103 (Panizzi *et al*., 2012). Based on previous biochemical studies showing CCDC103 is capable of stabilizing microtubules in solution (King and Patel-King, 2015), we hypothesized that these directed migration defects could be due to a loss of microtubule stability in CCDC103-deficient cells. Because increased microtubule instability is known to affect cell proliferation (Meyvisch *et al*., 1983), we first quantified the number of myeloid progenitors and differentiated macrophages and neutrophils in *smh* mutants using the *spi1b:EGFP, mpeg:YFP*, and *mpx:GFP* transgenes, respectively. Examining the number of each of these myeloid cells adjacent to the head and on the anterior yolks showed that *smh* mutants had fewer myeloid cells compared to their wild-type siblings (Fig. 3A-I). These observations were recapitulated when cells were quantified following whole-mount ISH (Fig. S5). Conversely, embryos injected with *ccdc103* mRNA had an increase in the numbers of *mpx+* and *mfap4+* cells compared to controls (Fig. S6). To determine if the decrease in these myeloid populations was due to reduced proliferation, we performed EdU pulse-chase experiments with *mpx:GFP* and *mpeg:YFP* embryos. Embryos were pulsed with EdU at the 20ss, fixed at 24 hours post-fertilization (hpf) and examined for EdU incorporation. We found that *smh* embryos displayed a reduced proportion of EdU^+^/*mpx:GFP*+ and EdU^+^/*mpeg:YFP*+ cells (Fig. 4A-F), indicating that the decreased numbers of anterior myeloid cells at 24 hpf was the result of a proliferation defect. Collectively, these results indicated that Ccdc103 promotes primitive myeloid proliferation in zebrafish embryos.

**Figure 3.**
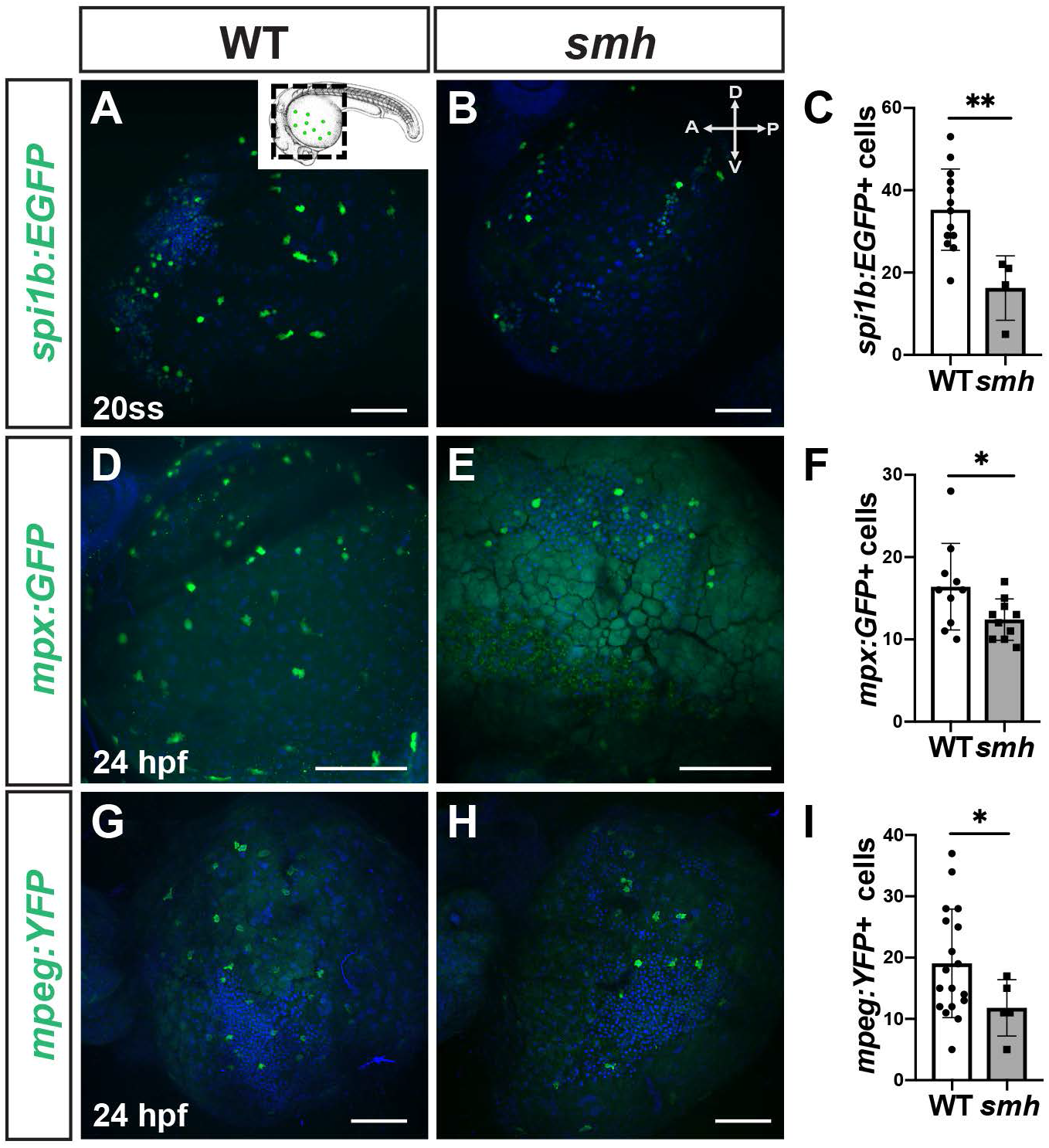
*smh* mutant embryos have fewer primitive myeloid cells at 24hpf. **(A**,**B)** Representative images of WT sibling (n=15) and *smh* embryos (n=5) carrying the *spi1b:EGFP* transgene at the 20ss. Inset indicates the region of the embryo being imaged. **(C)** Quantification of *spi1b:EGFP+* myeloid progenitors from one yolk hemisphere. (**D**,**E)** Representative images of WT (n=10) and *smh* (n=10) embryos with *mpx:GFP* transgene at 24hpf. **(F)** Quantification of *mpx:GFP+* cells from one yolk hemisphere. Each data point represents an individual embryo. **(G**,**H)** Representative images of WT sibling (n=19) and *smh* (n=9) embryos with *mpeg1*.*1:YFP* transgene at 24 hpf. **(I)** Quantification of *mpeg1*.*1:YFP+* myeloid progenitors from one yolk hemisphere. Scale bars: 100 μm.

**Figure 4.**
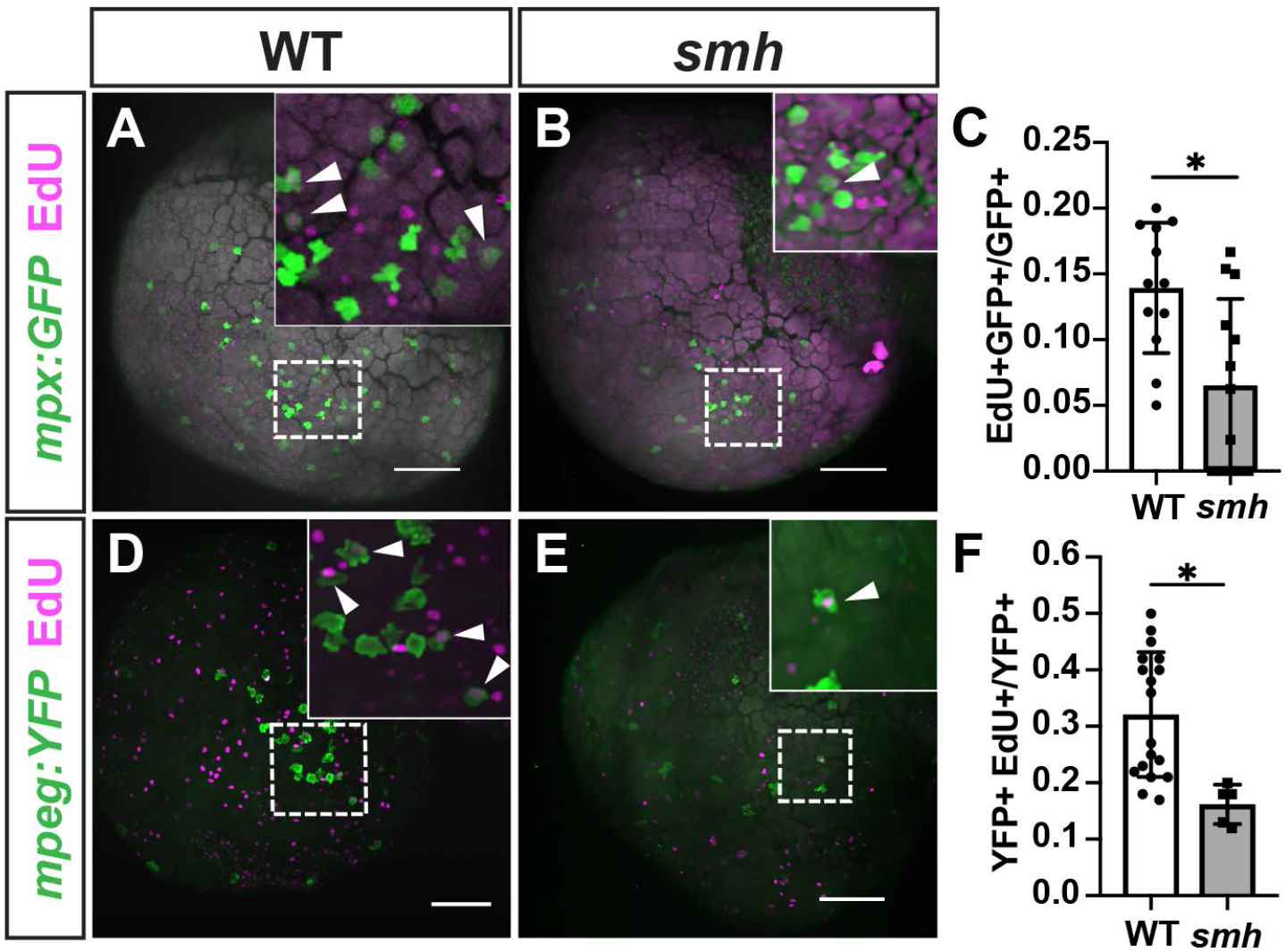
*Smh* myeloid cells are less proliferative. **(A**,**B)** Representative images of WT (n=12) and *smh* (n=19) embryos with *mpx:GFP* transgene pulsed with EdU at the 20ss and fixed at 24 hpf. White arrowheads indicate EdU+ cells. **(C)** Quantification of EdU+/*mpx:GFP+* cells from one yolk hemisphere, each data point representing an individual embryo. **(D**,**E)** Representative images of WT (n=20) and *smh* (n=9) embryos with *mpeg:YFP* transgene pulsed with EdU at the 20ss and fixed at 24 hpf. White arrowheads indicate EdU+ cells. **(F)** Quantification of EdU+/*mpeg:YFP+* cells from one yolk hemisphere, each data point representing an individual embryo. Scale bars: 100 μm.

### Myeloid cells in *smh* mutants have directed migration and morphology defects

Primary neutrophils isolated from PCD patients have impaired directed migration to chemokines in *in vitro* assays (Afzelius *et al*., 1980; Walter, Danielson and Reyes, 1990; Kantar *et al*., 1993; Cockx *et al*., 2017). Furthermore, microtubule destabilization via nocodazole treatment decreases the ability of neutrophils to migrate appropriately toward a chemical stimulus (Gundersen and Bulinski, 1988; Kadir *et al*., 2011; Yoo *et al*., 2012). To determine if neutrophils from *smh* mutant embryos have defects in directed migration, we generated sterile wounds on the yolks of zebrafish embryos and used time-lapse imaging to track the ability of stimulated neutrophils to home to the site of injury (Fig. 5A), as previously reported (Redd et al. 2006). At 24 hpf, neutrophils from wild-type (WT) sibling *mpx:GFP* embryos homed more effectively to wound sites than neutrophils from *smh* mutants (Fig. 5B-C; Video 1 and Video 2) and we calculated significantly decreased migration efficiency in both the X and Y dimensions in *smh* mutants compared to WT controls (Fig. 5D). Furthermore, *smh* neutrophils displayed a significantly decreased maximum track speed compared to controls (Fig. 5E). Similar results were obtained using *mpeg:YFP* embryos (Video 3, Video 4, and Fig. S7). In addition to improper migration, nocodazole-treated neutrophils display more spherical morphology than WT neutrophils (Yoo *et al*., 2010, 2012; Barros-Becker *et al*., 2017). Reminiscent of these studies, we also observed a higher mean sphericity index in neutrophils from *smh* mutants (Fig. 5F-H). Furthermore, high-resolution time-lapse imaging of individual neutrophils embryos showed that neutrophils from control *mpx:GFP* embryos predominantly adopt a clearly polarized elongated morphology, with defined uropods and lamellipodia characteristic of mature migrating neutrophils (Video 5 and Fig. 5I), while *smh*; *mpx:GFP* neutrophils predominantly lacked this characteristic shape and were rounded with minimal cytoplasmic extensions (Video 6 and Fig. 5J,K).

**Figure 5.**
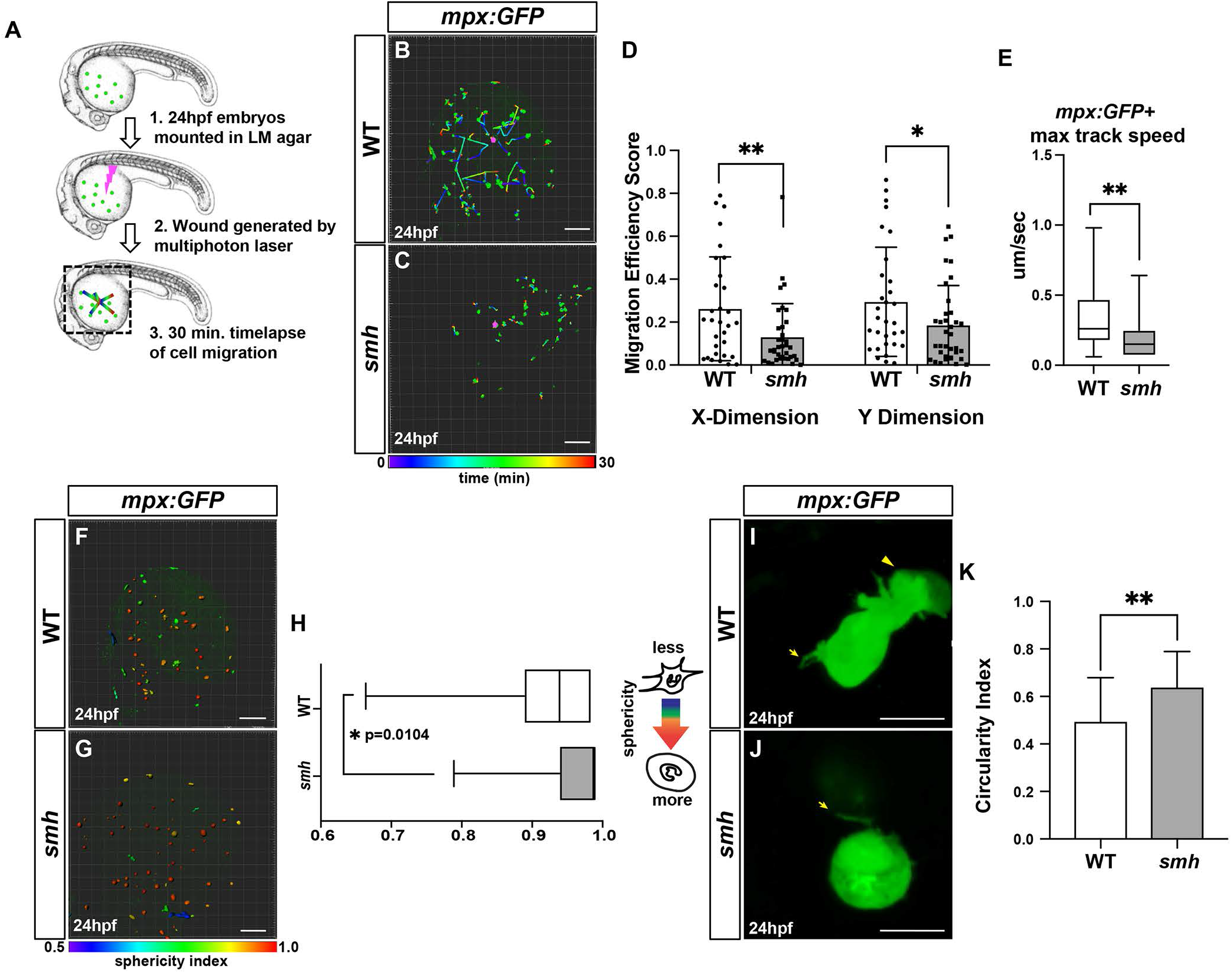
*Smh* mutant neutrophils display directed migration defects. **(A)** Schematic outlining wound generated with multiphoton laser. Dotted lines indicate the imaging area. **(B**,**C)** Representative confocal projection images of a wounded WT (n=6) and *smh* (n=6) embryos with the *mpx:gfp* transgene at 24hpf. Scale bars: 100 μm. **(D)** Quantification of migration efficiency scores calculated from point position data generated in Imaris. Each data point represents an individual cell from a minimum of 3 separate experiments, per genotype. **(E)** Quantification of the max maximum track speed of migrating cells. **(F**,**G)** Maximum confocal projections of wounded WT and *smh* embryos bearing the *mpx:gfp* transgene with individual cells projected as 3D surfaces and color-coded according to sphericity index as calculated by Imaris. Scale bars: 100 μm. **(H)** Quantification of the mean cell sphericity index. Box and whisker plots represent individual cells from all independent experiments at each time point imaged, in order to capture changes in cell sphericity over the course of the time-lapse sequence. **(I**,**J)** Representative single Z-slices from high-resolution live confocal imaging of individual *mpx:gfp*+ cells from WT (n=10) and *smh (n=12)* embryos. In I, arrow indicates uropod and arrowhead indicates lamellopodia. In J, arrow indicates cytoplasmic extension. Scale bars: 10 μm. **(K)** Mean circularity index for individually imaged cells as calculated in Imaris.

Given the correlation of myeloid defects in *smh* mutants to those in embryos with destabilized microtubules, we asked if the proliferation, directed migration and morphology defects in *smh* myeloid cells could be rescued by paclitaxel-mediated microtubule stabilization. A low dose of paclitaxel that did not cause overt developmental defects was administered to embryos at the 20ss followed by sterile wounding assays and EdU pulse-chase experiments. We observed that these low concentrations of paclitaxel were able to provide a partial rescue of the migration defects evident in *smh* embryos (Fig. 6A-C). Furthermore, paclitaxel treatment of *smh* embryos resulted in decreased cell sphericity compared to DMSO-treated *smh* embryos (Fig. 6E-G; Video 7 and Video 8). Additionally, paclitaxel administration rescued the proliferation defects in *smh* mutants (Fig. 6H). Taken together, our data support that Ccdc103-dependent microtubule stability is necessary to promote both myeloid cell proliferation and directed migration.

**Figure 6.**
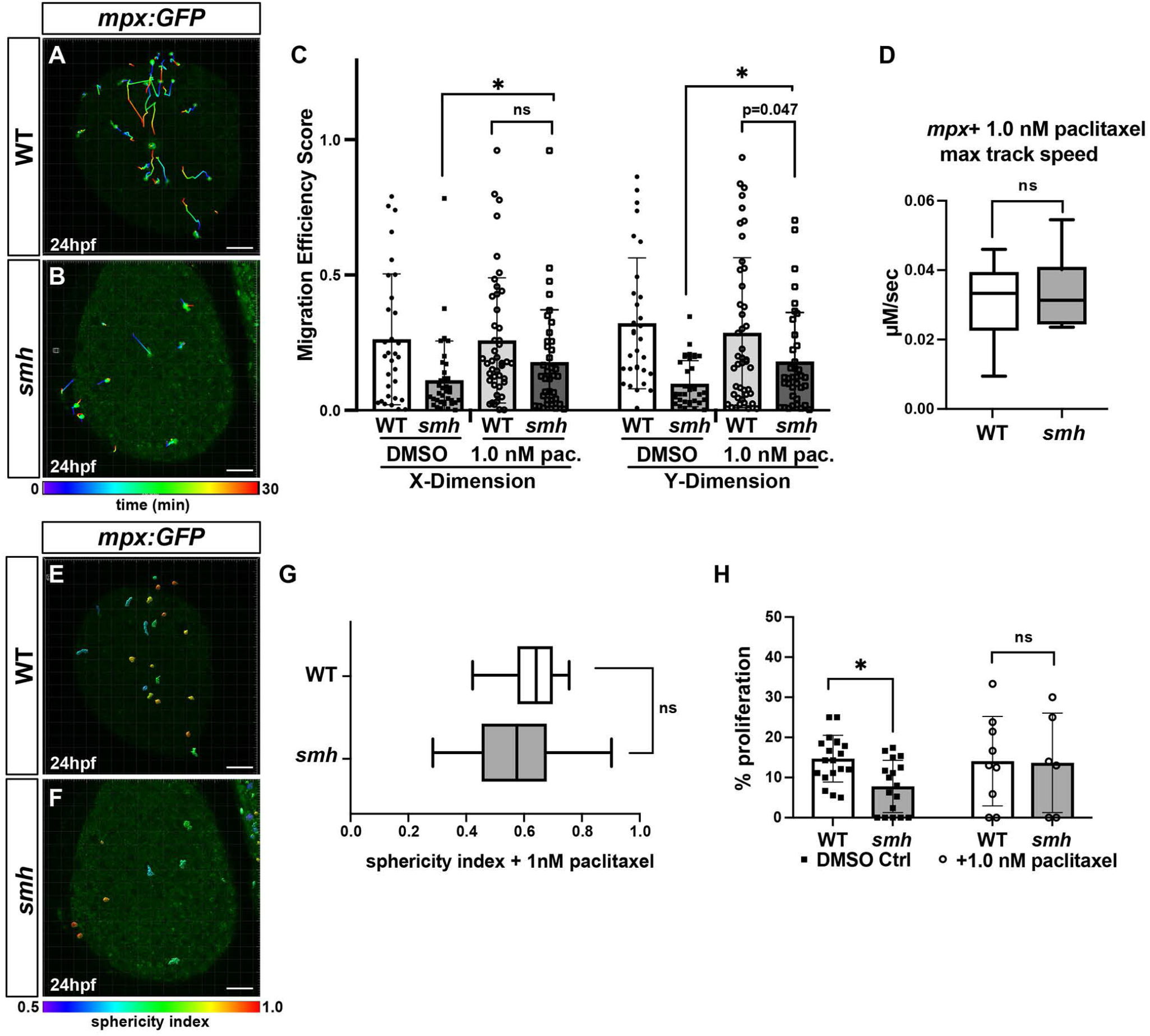
Paclitaxel can rescue proliferation and migration defects in *smh* mutants. **(A**,**B)** Representative confocal projection images from wounding experiments of paclitaxel-treated WT (n=4) and *smh* (n=4) *mpx:EGFP* transgenic embryos at 24 hpf. (**C**) Quantification of migration efficiency scores calculated from point position data generated in Imaris. Each data point represents an individual cell from a minimum of 3 separate experiments, per genotype, per treatment. **(D)** Quantification of max track speed. Each data point represents an individual cell. Cell tracks and sphericity were generated by Imaris. **(E**,**F)** Maximum confocal projections of wounded, paclitaxel-treated WT and *smh* embryos bearing the *mpx:gfp* transgene with individual cells projected as 3D surfaces and color-coded according to sphericity index as calculated by Imaris. Red - more spherical, blue - less spherical. **(G)** Mean cell sphericity indices as calculated in Imaris. (**H**) The percentage of EdU+/*mpx:EGFP*+ cells from transgenic WT and *smh* mutant embryos, pulsed with EdU at 17ss and treated with paclitaxel from 17ss to 24hpf. Scale bars: 100 μm.

### CCDC103 interacts with the microtubule-associated protein SPAG6

Based on biochemical evidence showing that CCDC103 forms self-organizing oligomers, others have hypothesized that CCDC103 may function as a molecular scaffold, anchoring other proteins at the microtubule and facilitating their function (King and Patel-King, 2020). However, very little is known about the network of CCDC103-interacting proteins. To elucidate conserved CCDC103-interacting proteins, which could provide mechanistic insights into how CCDC103 regulate microtubule stability within the cytoplasm of myeloid cells, we performed a yeast two-hybrid screen using the zebrafish Ccdc103 protein against a normalized human cDNA library. Interestingly, two of the interacting peptide sequences identified in this screen included the C-termini of DYNC1H1, validating the previous co-localization of CCDC103 and DYNC1H1 on microtubules that we observed with IHC (Fig. 2), and SPAG6 (Fig. 7A). The interaction with SPAG6 was particularly intriguing due to a number of phenotypic and functional similarities as CCDC103: mouse *Spag6* mutants display ciliary defects and many phenotypic hallmarks of PCD (Sapiro *et al*., 2002); Spag6 can promote effective migration of cortical neurons and proliferation and migration of mouse embryonic fibroblasts (Sapiro *et al*., 2002; Li, Mukherjee, Wu, Zhang, Maria E Teves, *et al*., 2015; Alciaturi *et al*., 2019) (Li, et al 2015) (Alciaturi *et al*., 2019); Spag6 has increased expression in myeloid leukemia lines (Cooley et al., 2016; Yang et al., 2015; Yin et al., 2018); and Spag6 can promote microtubule stability (Zheng *et al*., 2019). To validate these protein-protein interactions, we used the bioluminescent resonance transfer assay (BRET)-based LuTHy system (Trepte *et al*., 2018). The BRET assay confirmed CCDC103 interacts with both the C-terminal portion of DYNC1H1 (Fig. 7B) and the full-length SPAG6 (Fig. 7C). To determine whether this interaction required the presence of intact microtubules, we treated transfected cells with nocodazole and saw it significantly abrogated the interactions between CCDC103 and both DYNC1H1 and SPAG6 (Fig. 7B,C), while paclitaxel treatment did not affect the strength of the measured interactions.

**Figure 7.**
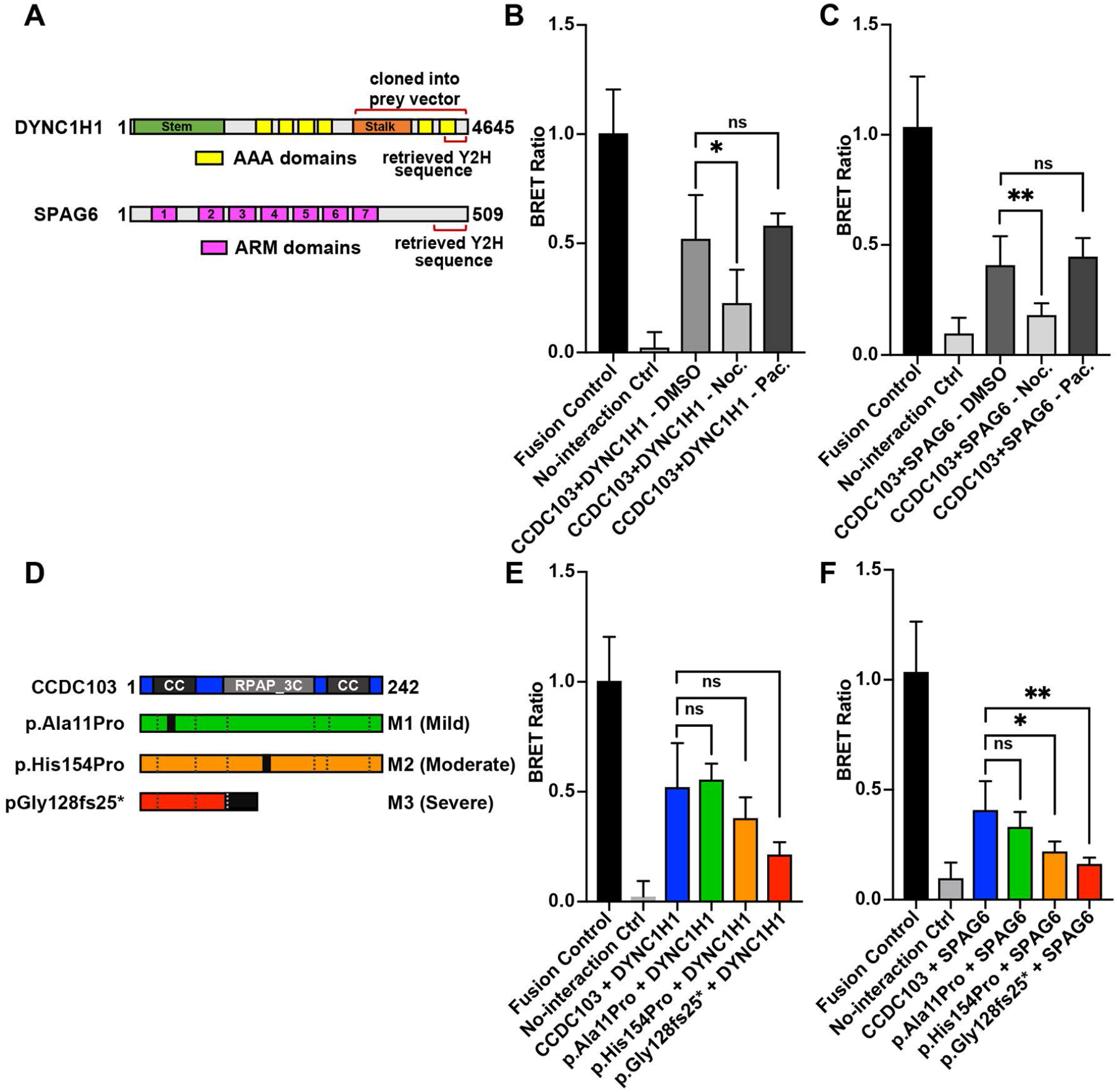
SPAG6 directly interacts with CCDC103. **(A)** Schematic of human SPAG6 and DYNC1H1 proteins. Specific domains identified by Y2H and portion of DYNC1H1 cloned into the LuTHy prey vector are indicated with brackets. The entirety of the *SPAG6* CDS was included in the respective LuTHy prey vector. **(B)** BRET ratios for the interaction between CCDC103 and DYNC1H1 in the presence of DMSO, nocodazole, and paclitaxel. **(C)** BRET ratios for the interaction between CCDC103 and SPAG6 in the presence of DMSO, nocodazole, and paclitaxel. **(D)** Schematic of WT CCDC103 protein and three mutations found in PCD patient mutations. Relative severity of the patient PCD phenotype associated with the mutation indicated in parentheses. **(E)** BRET ratios for the interaction between DYNC1H1 and WT and mutant CCDC103 proteins. **(F)** BRET ratios for the interaction between SPAG6 and WT and mutant CCDC103 proteins.

Currently, there are three known alleles in *CCDC103* that can cause PCD in humans, which result in highly variable clinical presentations (Fig. 7D) (Panizzi *et al*., 2012). To determine if these alleles affect the ability of CCDC103 to interact with DYNC1H1 and SPAG6, we performed the BRET assays with the *CCDC103* mutant alleles. Interestingly, the predicted severity of the mutations in CCDC103, which correlate with the overt severity of the patient PCD phenotypes, correlated with the loss of avidity in the interaction between CCDC103 and DYNC1H1 or SPAG6 (Fig. 7E,F) (Panizzi *et al*., 2012). Taken together, these results signify that patient mutations in CCDC103 that cause PCD disrupt microtubule-dependent interactions between CCDC103 and both DYNC1H1 and SPAG6.

### Loss of SPAG6 recapitulates *smh* mutant myeloid defects

In light of reports implicating SPAG6 in myeloid proliferation and cell migration (Sapiro *et al*., 2002; Jiang *et al*., 2019; Zheng *et al*., 2019), we interrogated the requirement of Spag6 in zebrafish myeloid cells. RT-PCR from flow-sorted *spi1b:EGFP* cells showed that zebrafish *spag6* is expressed in myeloid cells (Fig. S8). Zebrafish *spag6* mutants that harbor a 44 bp deletion predicted to cause a truncation via introduction of a premature stop codon in the second Armadillo repeat domain were generated using CRISPR/Cas9 (Fig. 8A and Fig. S8). Furthermore, this allele likely underwent non-sense mediated decay (Fig. S8). Fish carrying this *spag6* allele were homozygous viable and did not show significant overt signs of PCD that are observed in *smh* or other PCD zebrafish models, such as ventral curving of the body axis or pronephric cysts (Fig. S8) (Panizzi et al. 2012; Zariwala et al. 2013; Omori and Malicki 2006; Hjeij et al. 2013). However, *spag6* homozygous males were largely infertile, with fertilization rates of ∼13% (9 embryos fertilized out of 72 from one representative experiment) when crossed, consistent with observations of male factor infertility in *Spag6* mutant mice (Sapiro *et al*., 2002). Despite the lack of body curvature and pronephric defects in *spag6* mutants, we observed a low percentage of *spag6* mutants exhibited *situs inversus* of the heart (Fig. 8B-C), suggesting they may have hypomorphic motile cilia defects. With respect to myeloid cells, we found that *spag6* mutants have fewer myeloid progenitors (*spi1b:EGFP*^*+*^) (Fig. 8D,E,J), neutrophils (*mpx*^*+*^) (Fig. 8F,G,K), and macrophages (*mfap4*^*+*^) (Fig. 8H,L,I). Furthermore, neutrophils and macrophages in *spag6* mutants had decreased directed migration efficiency and more rounded morphology (Fig. 8M,N,O; Videos 9 and 10; Fig. S9). Thus, our data support that Spag6 regulates myeloid development and function in a similar manner to Ccdc103 that is consistent with promoting microtubule stability.

**Figure 8.**
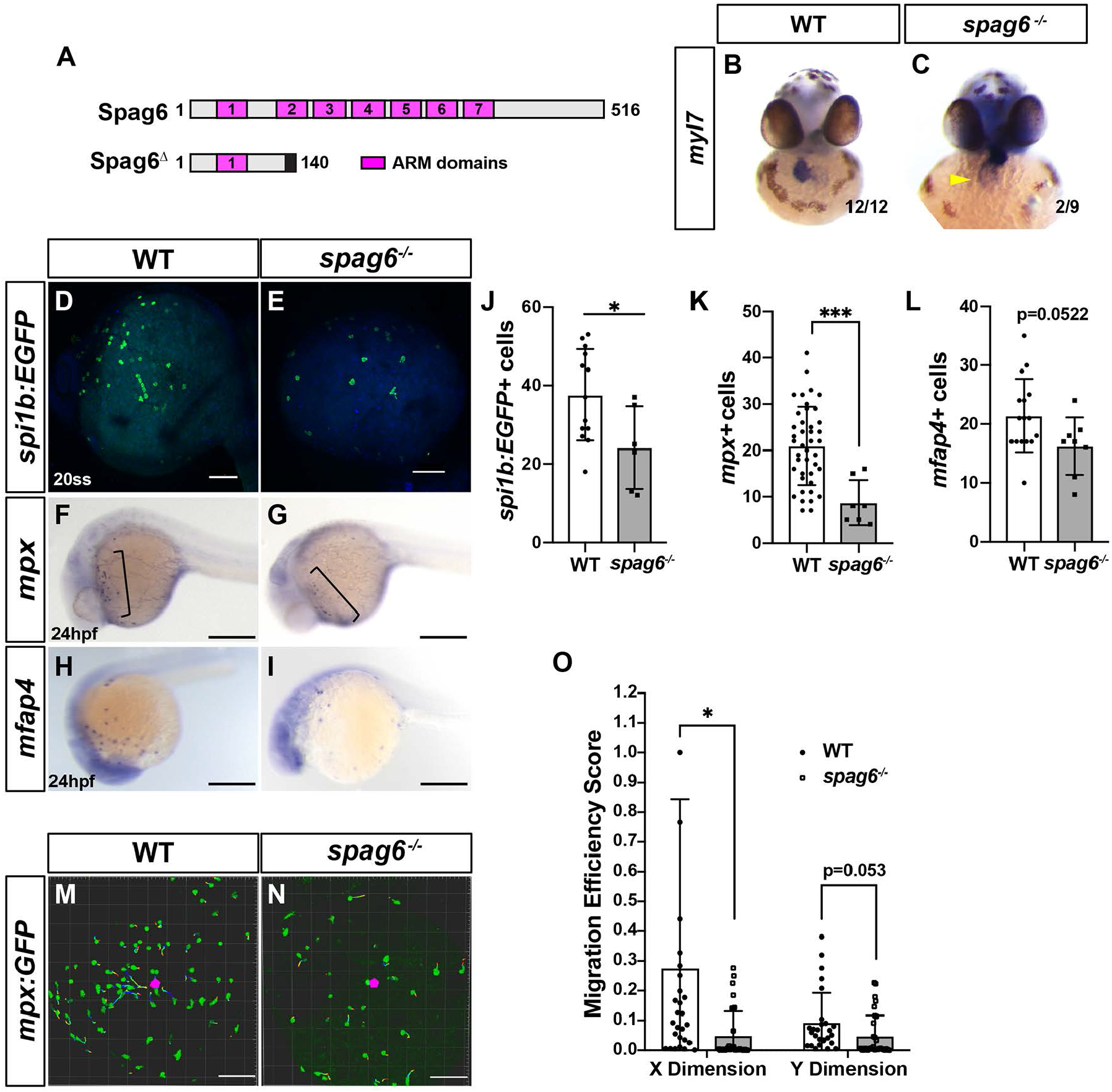
Spag6 is required for normal myeloid proliferation and migration. **(A)** Domain architecture for WT zebrafish Spag6 and the predicted truncation from the *spag6* mutant allele used. **(B**,**C)** ISH for *myl7* in WT and *spag6*^*-/-*^ embryos at 48hpf. *situs inversus* - yellow arrowhead in *spag6* mutant embryo. Fractions indicate the number of embryos with the given phenotype. **(D**,**E)** Whole mount IHC for *spi1b:EGFP* in WT (n=11) and *spag6* mutants (n=8). Scale bars: 100 μm. **(F**,**G)** Whole-mount ISH for the neutrophil marker *mpx* in WT (n=45) and *spag6* mutant (n=11) embryos at 24hpf. Scale bars: 300 μm. **(H**,**I)** Whole-mount ISH for the macrophage marker *mfap4* in WT (n=19) and *spag6* mutant (n=8) embryos at 24hpf. Scale bars: 300 μm. **(J-L)** Quantification of myeloid progenitors (*spi1b:EGFP+*), neutrophils (*mpx+*), and macrophages (*mfap4+*) from single yolk hemisphere of the embryos. **(M**,**N)** Representative confocal projection images from wounding experiments of WT sibling (n=4) and *spag6*^*-/-*^ (n=4) *mpx:EGFP* transgenic embryos at 24 hpf. Scale bars: 100 μm. **(O)** Migration efficiency scores from cell tracks of yolk wounding assays in WT and *spag6* mutant *mpx:EGFP+* embryos at 24 hpf.

## Discussion

Our data provide novel insights into cytoplasmic, extra-ciliary functions of CCDC103, in which mutations underlie a significant percentage of currently-identified PCD cases (Shoemark *et al*., 2018). Importantly, we show that CCDC103 has cilia-independent requirements regulating cytoplasmic microtubule stability that in turn promotes normal proliferation and migration of myeloid cells. Previous biochemical studies have shown that CCDC103 forms self-organizing oligomers, binds directly to and stabilizes cytoplasmic microtubules assembled in solution (King and Patel-King, 2015). Consequently, it was speculated that CCDC103 might serve as a scaffolding or adaptor protein, which could facilitate the function of multiple proteins that require localization at the microtubule. However, these proposed functions were not verified *in vivo* and it was unclear how they related to roles in the cilia. Our data corroborate the association of CCDC103 with microtubules *in vivo* and support that these interactions occur within the cytoplasm of myeloid cells. Intriguingly, while interactions between CCDC103 and axonemal dyneins have been demonstrated, we show that within the cytoplasm CCDC103 has microtubule-dependent interactions with cytoplasmic dynein (DYNC1H1). Thus, our results imply that one requirement of CCDC103 may be in stabilizing microtubule-dynein interactions to regulate cellular processes that require this cytoplasmic motor protein, including cargo transport and nuclear positioning (Vallee, McKenney and Ori-Mckenney, 2012; Roberts *et al*., 2013).

With the exceptions of axonemal dynein and microtubules, CCDC103-dependent protein complexes have not been identified (Werner, et al. 2012). Accordingly, in addition to DYNC1H1, we identified SPAG6 as a novel interactor of CCDC103. One reason it was an attractive candidate is because of the large body of literature indicating Spag6 promotes the same cellular processes as Ccdc103 in both ciliated and non-ciliated cells. Broadly, Spag6 has been shown to regulate cilia assembly, microtubule stability, proliferation, and migration (Li et al. 2015b; Zheng et al. 2019). While mutations in *SPAG6* have not as yet been identified as underlying cases of PCD in humans, *Spag6* KO mice have a higher incidence of neonatal death with ∼50% showing hydrocephalus (Sapiro *et al*., 2002). Furthermore, *Spag6* KO mice that survive have cilia defects in the tracheal epithelia and columnar cells of the middle ear, and the males are infertile (Sapiro *et al*., 2002). Like *ccdc103*, in zebrafish *spag6* is also expressed in the pronephros and spinal cord and is responsive to FoxJ1, a master regulator of the motile cilia program (Vij *et al*., 2012). Furthermore, a screen in zebrafish for genes that regulate motile cilia showed that morpholino oligo-mediated depletion of *spag6* resulted in pronephric cysts, but not other characteristics of PCD-associated defects in zebrafish, such as body curvature and hydrocephalus (Austin-Tse *et al*., 2013). While the *spag6* mutant allele we generated did show a modest increase in *situs inversus* of their hearts, which is indicative of motile cilia impairment, the mutants were viable and otherwise did not result in other overt PCD-related defects. However, similar to *Spag6* KO mice, the homozygous *spag6* zebrafish males were largely infertile. In mice, there is functional redundancy of Spag6 with Spag6-like (Spag6L) and *Spag6*; *Spag6L* KO mice have a higher frequency of early embryonic lethality and hydrocephalus (Cooley et al., 2016). However, zebrafish do not have a *spag6l* ortholog. Hence, the lack of other more severe overt phenotypes consistent with motile cilia defects could be due to redundancy with other *spag* genes. Although it has not been confirmed with zebrafish mutants, *spag1a* has been implicated in PCD in zebrafish (Knowles *et al*., 2013) as mutations in SPAG1 underlie PCD in humans (Knowles *et al*., 2013). Furthermore, we observe non-sense mediated decay with our *spag6* allele, which can trigger increased expression of related genes leading to genetic compensation and minimization of defects (Rossi *et al*., 2015).

In addition to the correlations in ciliated cells within the literature, our data support that zebrafish *smh* and *spag6* mutants both have previously unrecognized defects in myeloid proliferation and directed migration that are consistent with decreased microtubule stability. Increased SPAG6 expression is found in a broad range of myeloid cancers and myelodysplastic syndromes (Zheng et al. 2019; Steinbach et al. 2015; Yin et al. 2018b). Depletion of SPAG6 is associated with reduced proliferation of these leukemia lines, implying its increased expression contributes to their hyperplasia. Interestingly, it has also been suggested that SPAG6 promotes microtubule stability and longevity through promoting microtubule acetylation (Li, et al. 2015b), although the specific mechanisms by which this happens are not currently understood. While CCDC103 may be sufficient to increase microtubule stability independent of other factors *in vitro* (King and Patel-King 2015), it is feasible that SPAG6-mediated microtubule acetylation may contribute to CCDC103’s microtubule stabilizing effects *in vivo*. Moreover, it is interesting that CCDC103-SPAG6 interactions are more sensitive to the patient PCD-associated CCDC103 mutants than DYNC1H1, which implies the failure of these proteins to complex may be a key etiology driving both canonical PCD-associated and cilia-independent myeloid defects. Overall, we propose that CCDC103 may anchor the interactions of SPAG6 with microtubules to enhance their stability in the cytoplasm of myeloid cells, as well as within the axoneme of ciliated cells.

We currently do not understand if enhancement of microtubule stabilization in myeloid cells is a general requirement of canonical PCD-associated proteins or unique to CCDC103. Given that mutations in CCDC103 only account for a small number of known PCD cases (Knowles *et al*., 2013; Horani *et al*., 2016), we postulate that other PCD-associate proteins will have requirements promoting the production and function of myeloid cells through enhancing microtubule stability. All dynein axonemal assembly factors are also localized within the cytoplasm, supporting they have additional molecular functions in other cell types similar to Ccdc103. For example, Dyslexia susceptibility 1 candidate 1 (DYX1C1 - also called Dynein axonemal assembly factor 4 (DNAAF4)) is critical for motile ciliogenesis and axonemal dynein assembly and is intimately involved in neuronal migration (Casey et al. 2015; Tarkar et al. 2013; Chandrasekar et al. 2013). Furthermore, DYX1C1 has been shown to associate with cytoskeletal proteins, which may contribute to its effects on cellular migration (Tammimies *et al*., 2013). It has also been implicated in the proliferation of certain breast cancer cell subtypes because of its ability to modulate the function of the estrogen receptor (Massinen *et al*., 2009).

In conclusion, our study provides mechanistic insight into largely overlooked chemotactic defects found in neutrophils from PCD patients. Much remains to be learned about the pathophysiology of PCD. In particular, we envision that our observations may be relevant to the etiology of persistent pulmonary infections in these patients. Moving forward, deciphering the precise molecular mechanism by which CCD103-SPAG6 complexes promote microtubule stability and if there are broad requirements of PCD-associated proteins in facilitating myeloid proliferation and migration will enhance our understanding of immunological deficiency that may contribute to PCD, which could open avenues into novel treatment modalities to improve outcomes for these patients.

## Materials and Methods Ethics statement

All zebrafish husbandry and experiments were performed as described in approved IACUC protocols at the Cincinnati Children’s Hospital Medical Center.

### Zebrafish husbandry, transgenic and mutant lines

Adult zebrafish (Danio rerio) were raised and maintained under standard laboratory conditions (Westerfield, 2000). Zebrafish lines used were: *spi1b:EGFP*^*gl21Tg*^ (Ward *et al*., 2003), *mpeg1*.*1:YFP*^*w200Tg*^ (Roca and Ramakrishnan, 2013) ; *mpx:GFP*^*uwm1Tg*^ (Mathias *et al*., 2006); and *smh*^*tn222a*^ (Panizzi *et al*., 2012). *Smh* mutants were genotyped according to the protocols as previously described and with the primers listed (Table S1).

Zebrafish *spag6* mutants were created with standard CRISPR/Cas9 methods. Two guide RNAs (gRNAs) (Table S1) to exon 4 of *spag6* were injected with 150 pg each along with Cas9 protein (NEB, Cat. # M0646M). Injected embryos were screened for efficacy in creating deletions with PCR. The remaining F0 embryos were raised and subsequently screened for germline deletions in *spag6*. A *spag6* allele with a 44 bp deletion was used that is predicted to cause the protein to go out of frame after amino acid (AA) 131 and include a 9 AA extension prior to a stop codon. The allele used is designated *spag6*^*ci1013*^.

### mRNA injections

Zebrafish embryos were injected at the one cell stage with the following: 600 pg of *ccdc103* mRNA; 150 pg each of *scl* and *lmo2* mRNA; and 100 pg of *spi1b* mRNA. Capped mRNA was made using a Message Machine Kit (Ambion) according to manufacturer’s instructions.

### Whole-mount *in situ* hybridization

In situ hybridization (ISH) was performed as reported previously (D’Aniello *et al*., 2013). Probes for the following genes were used: *mpx* (ZDB-GENE-030131-9460*), mfap4 (*ZDB-GENE-040426-2246*), myl7 (*ZDB-GENE-991019-3) and *ccdc103 (*ZDB-GENE-040718-253). Embryos were visualized and photographed using a Zeiss M2BioV12 Stereo microscope.

### Directed migration and cell morphology analysis

*Smh* and *spag6* embryos bearing either the *mpeg1*.*1:YFP* or *mpx:GFP* transgenes were anaesthetized using ethyl 3-aminobenzoate methanesulfonate (tricaine) at a standard concentration of 164 mg/L (Sigma-Aldrich) and mounted in 0.6% low-melt agarose dissolved in embryo media. Wounds on the yolk were made in 24 hpf embryos with an upright Nikon FN1 microscope on an A1R confocal scanner. The scan area was decreased by 500x zoom, and the shutter was opened for 5 seconds. Wound sites were able to be visualized by autofluorescence in the GFP channel. Time-lapse images were captured with a 16x water immersion objective at 1.5X zoom. For each embryo, 150 micron Z-stacks were collected at 5 minute intervals for 1 hour (hr). Cell sphericity, track speed, and positional data were all captured and analyzed in Imaris (Bitplane).

Migration efficiency was calculated using Imaris by determining each individual cell’s motion vector in both the X and Y dimensions. These data were compared to a unique “ideal” vector for each cell, which represented the path that a given cell would need to take to reach the wound site by the conclusion of the time-lapse. Because the average distance in the Z dimension between a given cell and the wound site was only ∼10-15 microns, migration efficiency in that dimension was not calculated.

For imaging single cells, live wounded or unwounded embryos at either 24 hpf or 28 hpf (as specified) were imaged on a Nikon A1 using a 60X water-immersion objective at 5X zoom for the time-lapse duration and at the intervals specified. When indicated, embryos were treated with 1 nM paclitaxel (Sigma) solubilized in DMSO for 45 minutes prior to wounding and EdU assays. Experiments were then carried out as described above.

### Quantitative real time PCR

Total RNA isolation and real time quantitative PCR (RT-qPCR) was performed using previously reported methods (D’Aniello *et al*., 2013). Total RNA was isolated from staged embryos or FACS-sorted *spi1b:EGFP*^+^ myeloid progenitors that were homogenized in TRIzol (Ambion) and collected using RNeasy mini columns (Qiagen). The TURBO DNA-free kit (Applied Biosystems) was used to remove genomic contamination. 1 μg RNA was used for cDNA synthesis using the ThermoScript Reverse Transcriptase kit (Invitrogen). Quantitative real time PCR (qPCR) was performed using standard PCR conditions in a Bio-RadCFX PCR machine with Power SYBR Green PCR Master Mix (Applied Biosystems). Expression levels were standardized to *β*-actin or *GAPDH* expression. All experiments were performed in triplicate. cDNA for all myeloid cell lines, CD34+CD38-hematopoietic stem cells (HSCs) and whole cord blood was a gift of Dr. Jim Malloy.

### EdU assay

EdU labeling was carried out using the Click-iT EdU Alexa Fluor imaging kit (Molecular Probes) according to the manufacturer’s instructions. Briefly, dechorionated embryos at the 20ss were incubated with 10mM EdU for 30 minutes on ice, the EdU was washed out and embryos incubated at 28.5 degrees until 24 hpf. Embryos were then fixed overnight in 4% PFA in PBS at 24 hpf and the Click-iT reaction was performed according to the manufacturer’s protocol. After EdU incorporation, embryos were processed for immunohistochemistry using chicken anti-GFP antibody (1:250) and DAPI (1:5000). Embryos were imaged using a Nikon A1 confocal microscope and analyzed with Imaris (Bitplane).

### HL-60 Cell Culture and Differentiation

HL-60 cells were maintained in IMDM supplemented with 20% Bovine Growth Serum as per ATCC instructions. Cells were differentiated to neutrophils according to established methods (Tasseff *et al*., 2017). In short, neutrophil differentiation was performed by culturing for 96 hrs in 1 µM *all-*trans retinoic acid either (ATRA) in an 8-well chamber slide (Ibidi) or in suspension. HL-60 cells were differentiated into macrophage-like cells using 200ng/mL PMA for 48 hrs, in coated chamber slides or in suspension, according to established methods.

### Protein-protein Interaction Analysis

Ccdc103 was screened for candidate interacting proteins using the Matchmaker Gold Yeast Two-Hybrid System (Takara, Cat. # 630489) per the manufacturer’s instructions. Full length zebrafish *ccdc103* cDNA was used as bait and screened against a Mate & Plate – Universal Human Library (Normalized) (Takara, Cat. # 630481). 128 positive colonies were sequenced representing 66 different cDNAs. Candidate proteins were prioritized for additional analysis based on expression and known functions.

Bioluminescence-based resonance energy transfer (BRET) analysis of CCDC103-DYNC1H1 and CCDC103-SPAG6 interactions were carried out as described in Trepte et al (2018). HEK293T cells were plated at a density of 1×10^5 cells/mL in a 96-well plate and co-transfected with full-length CCDC103 CDS fused to a Nano-luciferase (NL)-Myc tag (Addgene plasmid # 113447) and either the full CDS (SPAG6) or in the case of DYNC1H1 a 1000 AA length of the C-terminal portion fused to a proteinA-mCitrine tag (Addgene plasmid # 113449). FuGENE® 6 Transfection Reagent (Promega) was used for all transfections. 48 hrs after transfection, Coelenterazine-h (Sigma-Aldrich Cat. # C3230) was added to a final concentration of 5 µM. Plates were incubated at 37° C for 15 minutes and then read on a Synergy H1 microplate reader (Biotek). Short-wavelength (SWL: 370-480 nM) and long-wavelength (LWL: 520-570 nM) luminescence values were collected on spectral scanning mode, with an integration time of 1,000 ms.

### Immunohistochemistry (IHC)

For IHC of differentiated and undifferentiated HL-60 cells, suspension cultures or chambered coverslips, the following protocol was used. First, growth media was aspirated from either coverslips or pelleted cells. Cells were washed 3X in room temperature PBS and fixed for 1 hr at 25°C in 4% PFA in PBS. Cells were washed 3X and then permeabilized in in PBS containing 0.1% Tween20/0.1% Triton X-100/1% DMSO for 15 minutes. Cells were washed again and then incubated with the specified primary antibodies (Table S2) for 1 hr at room temperature in a block solution (1% BSA, 1% DMSO, 0.1% Tween-20 in PBS), washed, and incubated for 1 hr in secondary antibody + blocking solution. Cells were protected from light. Following all washes, cells were mounted in SlowFade™ Diamond Antifade Mountant with DAPI (Thermofisher).

For all wholemount IHC, all of the above procedures were followed, except embryos were fixed at the developmental stages specified in 4% paraformaldehyde (PFA) in PBS either overnight at 4°C or for 2 hrs at 25°C (overnight) and all antibody incubation were performed overnight at 4°C. The affinity purified rabbit polyclonal pan-CCDC103 antibody was generated by YenZym Antibodies (www.yenzym.com) through a combination immunization approach with peptides for AAs #54-74 (SHLKPLEQKDKMGGKRFVPWN) of mouse Ccdc103 and AAs #54-74 (SHLKPLDRNDISGSPRKQPWN) of zebrafish Ccdc103. The antibody was verified via Western blot, comparison to a previously available commercial antibody (Abcam, Cat. # ab177558), and the lack of expression in *smh* mutants.

All imaging was performed on a Nikon Eclipse Ti inverted microscope on a Nikon A1R confocal and processed in NIS-Elements AR or Imaris, as appropriate. Images were processed using Nikon denoise.ai functionality.

### Statistical analysis

Unless otherwise specified, all statistical tests were carried out using two-tailed Student’s t-test with an *α* value of p<0.05. All statistical analyses were performed in GraphPad Prism 6.0. All data presented in this paper is given as the pooled results of at least three independent experiments with embryos gathered from independent crosses.

## Acknowledgements

We thank members of the Waxman lab for helpful discussions.

## Competing interests

The authors have no competing interests.

## Figure Legends

**Video 1. WT - *mpx:GFP* wounding assay**.

**Video 2. *smh* mutant - *mpx:GFP* wounding assay**.

**Video 3. WT - *mpx:GFP+* cell @ 60X**.

**Video 4. *smh* mutant - *mpx:GFP+* cell @ 60X**.

**Video 5. WT - *mpeg:GFP* wounding assay**.

**Video 6. *smh* mutant - *mpeg:GFP* wounding assay**.

**Video 7. WT - *mpx:GFP* + paclitaxel in wounding assay**.

**Video 8. *smh* - *mpx:GFP* + paclitaxel in wounding assay**.

**Video 9. WT (*spag6*)- *mpx:GFP* wounding assay**.

**Video 10. *spag6* mutant - *mpx:GFP* wounding assay**.

## References

Afzelius, B. A. et al. (1980) ‘Structure and function of neutrophil leukocytes from patients with the immotile-cilia syndrome.’, Acta medica Scandinavica, 208(3), pp. 145–54. doi: 10.1111/j.0954-6820.1980.tb01169.x.

Alciaturi, J. et al. (2019) ‘Distribution of sperm antigen 6 (SPAG6) and 16 (SPAG16) in mouse ciliated and non-ciliated tissues’, Journal of Molecular Histology, 50(3), pp. 189–202. doi: 10.1007/s10735-019-09817-z.

Austin-Tse, C. et al. (2013) ‘Zebrafish Ciliopathy Screen Plus Human Mutational Analysis Identifies C21orf59 and CCDC65 Defects as Causing Primary Ciliary Dyskinesia’, The American Journal of Human Genetics, 93(4), pp. 672–686. doi: 10.1016/j.ajhg.2013.08.015.

Barros-Becker, F. et al. (2017) ‘Live imaging reveals distinct modes of neutrophil and macrophage migration within interstitial tissues.’, Journal of cell science, 130(22), pp. 3801–3808. doi: 10.1242/jcs.206128.

Breitman, T. R., Selonick, S. E. and Collins, S. J. (1980) ‘Induction of differentiation of the human promyelocytic leukemia cell line (HL-60) by retinoic acid.’, Proceedings of the National Academy of Sciences of the United States of America, 77(5), pp. 2936–40. doi: 10.1073/pnas.77.5.2936.

Casey, J. P. et al. (2015) ‘A case report of primary ciliary dyskinesia, laterality defects and developmental delay caused by the co-existence of a single gene and chromosome disorder’, BMC Medical Genetics, 16(1), p. 45. doi: 10.1186/s12881-015-0192-z.

Chandrasekar, G. et al. (2013) ‘The Zebrafish Orthologue of the Dyslexia Candidate Gene DYX1C1 Is Essential for Cilia Growth and Function’, PLoS ONE. doi: 10.1371/journal.pone.0063123.

Cockx, M. et al. (2017) ‘Neutrophils from Patients with Primary Ciliary Dyskinesia Display Reduced Chemotaxis to CXCR2 Ligands.’, Frontiers in immunology, 8, p. 1126. doi: 10.3389/fimmu.2017.01126.

Cockx, M. et al. (2018) ‘Chemoattractants and cytokines in primary ciliary dyskinesia and cystic fibrosis: key players in chronic respiratory diseases’, Cellular & Molecular Immunology, 15(4), pp. 312–323. doi: 10.1038/cmi.2017.118.

Cooley, L. F., El Shikh, M. E., Li, W., Keim, R. C., Zhang, Z., et al. (2016) ‘Impaired immunological synapse in sperm associated antigen 6 (SPAG6) deficient mice’, Scientific Reports, 6(1), p. 25840. doi: 10.1038/srep25840.

Cooley, L. F., El Shikh, M. E., Li, W., Keim, R. C., Zhang, Z. Z., et al. (2016) ‘No Title’, 6(1), p. 25840. doi: 10.1038/srep25840.

D’Aniello, E. et al. (2013) ‘Depletion of retinoic acid receptors initiates a novel positive feedback mechanism that promotes teratogenic increases in retinoic acid.’, PLoS genetics, 9(8), p. e1003689. doi: 10.1371/journal.pgen.1003689.

Damseh, N. et al. (2017) ‘Primary ciliary dyskinesia: mechanisms and management.’, The application of clinical genetics, 10, pp. 67–74. doi: 10.2147/TACG.S127129.

Englander, L. L. and Malech, H. L. (1981) ‘Abnormal movement of polymorphonuclear neutrophils in the Immotile Cilia Syndrome. Cinemicrographic analysis.’, Experimental cell research, 135(2), pp. 468–72. Available at: http://www.ncbi.nlm.nih.gov/pubmed/6975720 (Accessed: 5 September 2018).

Finetti, F. et al. (2009) ‘Intraflagellar transport is required for polarized recycling of the TCR/ CD3 complex to the immune synapse’, Nature Cell Biology. doi: 10.1038/ncb1977.

Gering, M. et al. (2003) ‘Lmo2 and Scl/Tal1 convert non-axial mesoderm into haemangioblasts which differentiate into endothelial cells in the absence of Gata1.’, Development (Cambridge, England), 130(25), pp. 6187–99. doi: 10.1242/dev.00875.

Gundersen, G. G. and Bulinski, J. C. (1988) ‘Selective stabilization of microtubules oriented toward the direction of cell migration.’, Proceedings of the National Academy of Sciences of the United States of America, 85(16), pp. 5946–50. doi: 10.1073/pnas.85.16.5946.

Hjeij, R. et al. (2013) ‘ARMC4 mutations cause primary ciliary dyskinesia with randomization of left/right body asymmetry’, American Journal of Human Genetics. doi: 10.1016/j.ajhg.2013.06.009.

Horani, A. et al. (2016) ‘Genetics and biology of primary ciliary dyskinesia’, Paediatric Respiratory Reviews. W.B. Saunders Ltd, pp. 18–24. doi: 10.1016/j.prrv.2015.09.001.

Jiang, M. et al. (2019) ‘Upregulation of SPAG6 in Myelodysplastic Syndrome: Knockdown Inhibits Cell Proliferation via AKT/FOXO Signaling Pathway.’, DNA and cell biology, 38(5), pp. 476–484. doi: 10.1089/dna.2018.4521.

Kadir, S. et al. (2011) ‘Microtubule remodelling is required for the front-rear polarity switch during contact inhibition of locomotion’, Journal of Cell Science, 124(15), pp. 2642–2653. doi: 10.1242/jcs.087965.

Kantar, A. et al. (1993) ‘Membrane Fluidity of Polymorphonuclear Leukocytes from Children with Primary Ciliary Dyskinesia’, Pediatric Research, 34(6), pp. 725–728. doi: 10.1203/00006450-199312000-00006.

King, S. M. and Patel-King, R. S. (2015) ‘The oligomeric outer dynein arm assembly factor CCDC103 is tightly integrated within the ciliary axoneme and exhibits periodic binding to microtubules’, Journal of Biological Chemistry, 290(12), pp. 7388–7401. doi: 10.1074/jbc.M114.616425.

King, S. M. and Patel-King, R. S. (2020) ‘The outer dynein arm assembly factor CCDC103 forms molecular scaffolds through multiple self-interaction sites’, Cytoskeleton, 77(1–2), pp. 25–35. doi: 10.1002/cm.21591.

Knowles, M. R. et al. (2013) ‘Mutations in SPAG1 Cause Primary Ciliary Dyskinesia Associated with Defective Outer and Inner Dynein Arms’, The American Journal of Human Genetics, 93(4), pp. 711–720. doi: 10.1016/j.ajhg.2013.07.025.

Leigh, M. W. et al. (2009) ‘Clinical and genetic aspects of primary ciliary dyskinesia/Kartagener syndrome’, Genetics in Medicine, 11(7), pp. 473–487. doi: 10.1097/GIM.0b013e3181a53562.

Li, W., Mukherjee, A., Wu, J., Zhang, L., Teves, Maria E, et al. (2015) ‘Sperm Associated Antigen 6 (SPAG6) Regulates Fibroblast Cell Growth, Morphology, Migration and Ciliogenesis.’, Scientific reports, 5, p. 16506. doi: 10.1038/srep16506.

Li, W., Mukherjee, A., Wu, J., Zhang, L., Teves, Maria E., et al. (2015a) ‘Sperm Associated Antigen 6 (SPAG6) Regulates Fibroblast Cell Growth, Morphology, Migration and Ciliogenesis’, Scientific Reports, 5(1), p. 16506. doi: 10.1038/srep16506.

Li, W., Mukherjee, A., Wu, J., Zhang, L. Teves, Maria E., et al. (2015b) ‘Sperm Associated Antigen 6 (SPAG6) Regulates Fibroblast Cell Growth, Morphology, Migration and Ciliogenesis’, Scientific Reports, 5(1), p. 16506. doi: 10.1038/srep16506.

Massinen, S. et al. (2009) ‘Functional interaction of DYX1C1 with estrogen receptors suggests involvement of hormonal pathways in dyslexia’, Human Molecular Genetics. doi: 10.1093/hmg/ddp215.

Mathias, J. R. et al. (2006) ‘Resolution of inflammation by retrograde chemotaxis of neutrophils in transgenic zebrafish’, Journal of Leukocyte Biology, 80(6), pp. 1281–1288. doi: 10.1189/jlb.0506346.

McClure-Begley, T. D. and Klymkowsky, M. W. (2017) ‘Nuclear roles for cilia-associated proteins’, Cilia. doi: 10.1186/s13630-017-0052-x.

Meyvisch, C. et al. (1983) ‘Invasiveness and tumorigenicity of MO4 mouse fibrosarcoma cells pretreated with microtubule inhibitors’, Clinical & Experimental Metastasis. doi: 10.1007/BF00118469.

Millius, A. and Weiner, O. D. (2010) ‘Manipulation of Neutrophil-Like HL-60 Cells for the Study of Directed Cell Migration’, Methods in molecular biology (Clifton, N.J.), 591, p. 147. doi: 10.1007/978-1-60761-404-3_9.

Mj, R. et al. (2006) ‘Imaging Macrophage Chemotaxis in Vivo: Studies of Microtubule Function in Zebrafish Wound Inflammation’, Cell motility and the cytoskeleton, 63(7). doi: 10.1002/CM.20133.

Omori, Y. and Malicki, J. (2006) ‘oko meduzy and Related crumbs Genes Are Determinants of Apical Cell Features in the Vertebrate Embryo’, Current Biology, 16(10), pp. 945–957. doi: 10.1016/j.cub.2006.03.058.

Panizzi, J. R. et al. (2012) ‘CCDC103 mutations cause primary ciliary dyskinesia by disrupting assembly of ciliary dynein arms’, Nature Genetics, 44(6), pp. 714–719. doi: 10.1038/ng.2277.

Roberts, A. J. et al. (2013) ‘Functions and mechanics of dynein motor proteins’, Nature Reviews Molecular Cell Biology. doi: 10.1038/nrm3667.

Roca, F. J. and Ramakrishnan, L. (2013) ‘TNF dually mediates resistance and susceptibility to mycobacteria via mitochondrial reactive oxygen species’, Cell, 153(3), pp. 521–534. doi: 10.1016/j.cell.2013.03.022.

Rossi, A. et al. (2015) ‘Genetic compensation induced by deleterious mutations but not gene knockdowns’, Nature. doi: 10.1038/nature14580.

Sapiro, R. et al. (2002) ‘Male infertility, impaired sperm motility, and hydrocephalus in mice deficient in sperm-associated antigen 6.’, Molecular and cellular biology, 22(17), pp. 6298–305. doi: 10.1128/mcb.22.17.6298-6305.2002.

Shapiro, A. J. et al. (2018) ‘Diagnosis of Primary Ciliary Dyskinesia. An Official American Thoracic Society Clinical Practice Guideline’, American Journal of Respiratory and Critical Care Medicine, 197(12), pp. e24–e39. doi: 10.1164/rccm.201805-0819ST.

Shoemark, A. et al. (2018) ‘High prevalence of CCDC103 p.His154Pro mutation causing primary ciliary dyskinesia disrupts protein oligomerisation and is associated with normal diagnostic investigations’, Thorax, 73(2), pp. 157–166. doi: 10.1136/thoraxjnl-2017-209999.

Steinbach, D. et al. (2015) ‘Prospective Validation of a New Method of Monitoring Minimal Residual Disease in Childhood Acute Myelogenous Leukemia’, Clinical Cancer Research, 21(6), pp. 1353–1359. doi: 10.1158/1078-0432.CCR-14-1999.

Tammimies, K. et al. (2013) ‘Molecular networks of DYX1C1 gene show connection to neuronal migration genes and cytoskeletal proteins’, Biological Psychiatry. doi: 10.1016/j.biopsych.2012.08.012.

Tarkar, A. et al. (2013) ‘DYX1C1 is required for axonemal dynein assembly and ciliary motility’, Nature Genetics. doi: 10.1038/ng.2707.

Tasseff, R. et al. (2017) ‘An Effective Model of the Retinoic Acid Induced HL-60 Differentiation Program.’, Scientific reports, 7(1), p. 14327. doi: 10.1038/s41598-017-14523-5.

Trepte, P. et al. (2018) ‘LuTHy: a double-readout bioluminescence-based two-hybrid technology for quantitative mapping of protein-protein interactions in mammalian cells.’, Molecular systems biology, 14(7), p. e8071. doi: 10.15252/msb.20178071.

Valerius, N. H., Knudsen, B. B. and Pedersen, M. (1983) ‘Defective neutrophil motility in patients with primary ciliary dyskinesia’, European Journal of Clinical Investigation, 13(6), pp. 489–494. doi: 10.1111/j.1365-2362.1983.tb00134.x.

Vallee, R. B., McKenney, R. J. and Ori-Mckenney, K. M. (2012) ‘Multiple modes of cytoplasmic dynein regulation’, Nature Cell Biology. doi: 10.1038/ncb2420.

Vij, S. et al. (2012) ‘Evolutionarily Ancient Association of the FoxJ1 Transcription Factor with the Motile Ciliogenic Program’, PLoS Genetics. doi: 10.1371/journal.pgen.1003019.

Walter, R. J., Danielson, J. R. and Reyes, H. M. (1990) ‘Characterization of a chemotactic defect in patients with Kartagener syndrome.’, Archives of otolaryngology--head & neck surgery, 116(4), pp. 465–9. doi: 10.1001/archotol.1990.01870040087020.

Ward, A. C. et al. (2003) ‘The zebrafish spi1 promoter drives myeloid-specific expression in stable transgenic fish’, Blood, 102(9), pp. 3238–3240. doi: 10.1182/blood-2003-03-0966.

Werner, C. et al. (2012) ‘CCDC103 encodes a novel cilia dynein arm factor that is mutated in primary ciliary dyskinesia’, European Respiratory Journal.

Yang, B. et al. (2015) ‘SPAG6 silencing inhibits the growth of the maligannt myeloid cell lines SKM-1 and K562 via activating p53 and caspase activation-dependent apoptosis’, International Journal of Oncology, 46(2), pp. 649–656. doi: 10.3892/ijo.2014.2768.

Yin, J. et al. (2018a) ‘SPAG6 silencing induces apoptosis in the myelodysplastic syndrome cell line SKM-1 via the PTEN/PI3K/AKT signaling pathway in vitro and in vivo.’, International journal of oncology, 53(1), pp. 297–306. doi: 10.3892/ijo.2018.4390.

Yin, J. et al. (2018b) ‘SPAG6 silencing induces apoptosis in the myelodysplastic syndrome cell line SKM-1 via the PTEN/PI3K/AKT signaling pathway in vitro and in vivo.’, International journal of oncology, 53(1), pp. 297–306. doi: 10.3892/ijo.2018.4390.

Yoo, S. K. et al. (2010) ‘Differential Regulation of Protrusion and Polarity by PI(3)K during Neutrophil Motility in Live Zebrafish’, Developmental Cell, 18(2), pp. 226–236. doi: 10.1016/j.devcel.2009.11.015.

Yoo, S. K. et al. (2012) ‘The role of microtubules in neutrophil polarity and migration in live zebrafish.’, Journal of cell science, 125(Pt 23), pp. 5702–10. doi: 10.1242/jcs.108324.

Yuan, S. and Sun, Z. (2013) ‘Expanding horizons: Ciliary proteins reach beyond cilia’, Annual Review of Genetics. doi: 10.1146/annurev-genet-111212-133243.

Zariwala, M. A. et al. (2013) ‘ZMYND10 is mutated in primary ciliary dyskinesia and interacts with LRRC6’, American Journal of Human Genetics. doi: 10.1016/j.ajhg.2013.06.007.

Zheng, D.-F. et al. (2019) ‘The Emerging Role of Sperm-Associated Antigen 6 Gene in the Microtubule Function of Cells and Cancer.’, Molecular therapy oncolytics, 15, pp. 101–107. doi: 10.1016/j.omto.2019.08.011.

